# Mate Competition Drives Aggressive Behaviour in Female *Drosophila*

**DOI:** 10.1101/2022.02.07.479369

**Authors:** Miguel Gaspar, Sophie Dias, Maria Luísa Vasconcelos

**Affiliations:** Champalimaud Research, Champalimaud Foundation, Lisbon 1400-038, Portugal

**Keywords:** *Drosophila melanogaster*, female aggression, mate competition, mating drive, olfaction, OR47b

## Abstract

Aggression is an adaptive set of behaviours that allows animals to compete against one another in an environment of limited resources. In *Drosophila* such aggressive behaviour has been extensively studied in males. Despite recent work highlighting territorial defence in females, female aggression in *Drosophila* is still poorly understood. Indeed, whether females compete for mating partners, as males do, has remained unknown so far. In the present work, we report that *Drosophila melanogaster* females reliably display aggression towards mating pairs. This aggressive behaviour is positively associated with the female’s mating drive and relies heavily on olfaction. Furthermore, we found that food odour in combination with OR47b-dependent fly odour sensing are required for proper expression of aggressive behaviour. Taken together, we describe a social context linked to reproduction in which *Drosophila* females aspiring to mate produce consistent and stereotyped displays of aggression. These findings open the door for further inquiries into the neural mechanisms that govern this behaviour.

## INTRODUCTION

Intraspecific aggression often emerges in the context of competition for resources. Typically, males fight for mates and food while females fight for food and nest sites^1^. These behavioural contexts have been used extensively in the studies of aggression in the fruit fly, which has been an important model system to study this behaviour, since it displays high levels of complex and highly structured aggression in the presence of a food patch and conspecifics^2,3^. Importantly, since the fly is a powerful genetic model system, the study of fly aggressive behaviour has been accompanied by an impressive body of work dissecting the neuronal circuits underlying aggression using, as a starting point, neurons that express the sexual determination genes *fruitless* and *doublesex*^4–23,23– 27^. Furthermore, unlike male aggression, female aggression does not lead to the establishment of dominance relationships between contest winners and losers^22,28^. The cues that promote aggression also differ between the sexes. The presence of a decapitated female in the food patch will increase fighting intensity in males^2^ whereas females will fight more vigorously in the presence of live yeast^29^. In males the perception of male-specific pheromones, both volatile and non-volatile, as well as the sound of conspecifics have been shown to regulate aggression^21,23,30^. Analogous signals in females have not yet been identified. Another interesting feature of female aggressive behaviour is that it is strongly stimulated by mating^31,32^, which is consistent with the notion that females fight for resources to improve the odds of their offspring. However, in certain circumstances females of other species compete for mates^33–36^. Female competition for males may arise whenever the adult sex ratio of the population becomes biased towards females, when there is a high degree of reproductive synchrony, or in mating systems where females mate with multiple partners^37,38^. Here we address competition between females for a mating partner in the fruit fly *Drosophila melanogaster*. We analysed the interactions between two sexually receptive females and a male, i.e., an adult sex ratio biased towards females. We found that, although females did not appear to compete during courtship, they did become aggressive once a pair copulated. We explored how reproductive parameters are affected by aggression and found no clear effect on any of the measured parameters. We show that the amount of aggression displayed in these circumstances has a positive correlation with the female mating drive, which is the opposite of what had previously been reported for food patch competition^31,32^. Finally, we show that food odour is a pre-requisite for these aggressive displays and that conspecific sensing through the odorant receptor OR47b contributes substantially to generate aggressive displays.

Our results reveal a new context of female aggression in *Drosophila melanogaster*, widening our understanding of how behaviour is modulated by different internal states and external circumstances.

## RESULTS

### Female aggressive behaviour is reliable and stereotyped in the presence of a mating pair

To investigate whether females compete for males, we started by pairing naive males with two virgin females and recorded their interactions for one hour. This allows us to capture both the entire courtship period leading up to copulation, as well as the entirety of copulation from start to finish. We define an aggression bout as any period of time where females display consecutive headbutts and/or shoving, the highest intensity aggressive behaviours in females^22,29,39^ (see Movie S1). While we failed to observe any kind of agonistic interaction between the animals while courtship is ongoing, we reliably found that a lot of their encounters (Figure S1A) resulted in female aggressive behaviour during copulation (Figures 1A, 1B, 1C, and S1B). We then decided to characterize when and how female flies display this behaviour. We started by investigating whether aggressive behaviour is displayed at random during copulation. If this were so, we would expect to find its occurrence to be uniformly distributed over copulation time. We found, however, that aggression is mostly concentrated in the first half of copulation (Figure 1D), around 5 to 9 minutes (Figure S1C). Given that ejaculation is expected to occur within that period of time^40^, these findings are in line with a possible strategy to dismount the copulating male; alternatively, female aggression drive could decay drastically after the initial bouts. Once copulation ends, a new phase of courtship with the second female follows, which can culminate in copulation with that second female. However, no aggression was ever observed after the end of the first copulation, either during courtship of the second female, or during the second copulation (data not shown). Next, we wondered whether aggressive females showed any preference in targeting either the mating male or the mating female. When identifying the target of each aggressive bout, we found that they are more or less evenly distributed between both sexes (Figure 1E). It thus seems that despite the high levels of aggressive displays and its focus on early copulation, females target both mating individuals indiscriminately. Next, we used positional information gathered from tracking data to further characterize female behaviour during aggressive bouts. We found that females perform aggressive displays with both facing angles and angles between the flies comprised within a narrow range: the former falling between 0º and 30º (Figure 1F), and the latter between 60º and 120º (Figure 1G). These results reveal that rather than head-to-head attacks, as is typically observed in male-male aggression^2^, females attack the flanks of mating pairs. Moreover, we found that although females will headbutt and shove either mating fly (Figure 1E), they preferentially target the posterior half of the female (Figure 1H), an area that, corresponds to where the male is located on top of the mating female. Whether this specific targeting is an adaptive strategy employed by the female to affect the reproductive outcome of the ongoing copulation, remains unclear. Taken together, we have shown that females reliably fight during copulation of a mating pair in a stereotypical fashion, flanking the posterior half of the mating female (Figure 1I). We next investigated whether these aggressive displays have any bearing in the reproductive output of either female.

**Figure 1.**
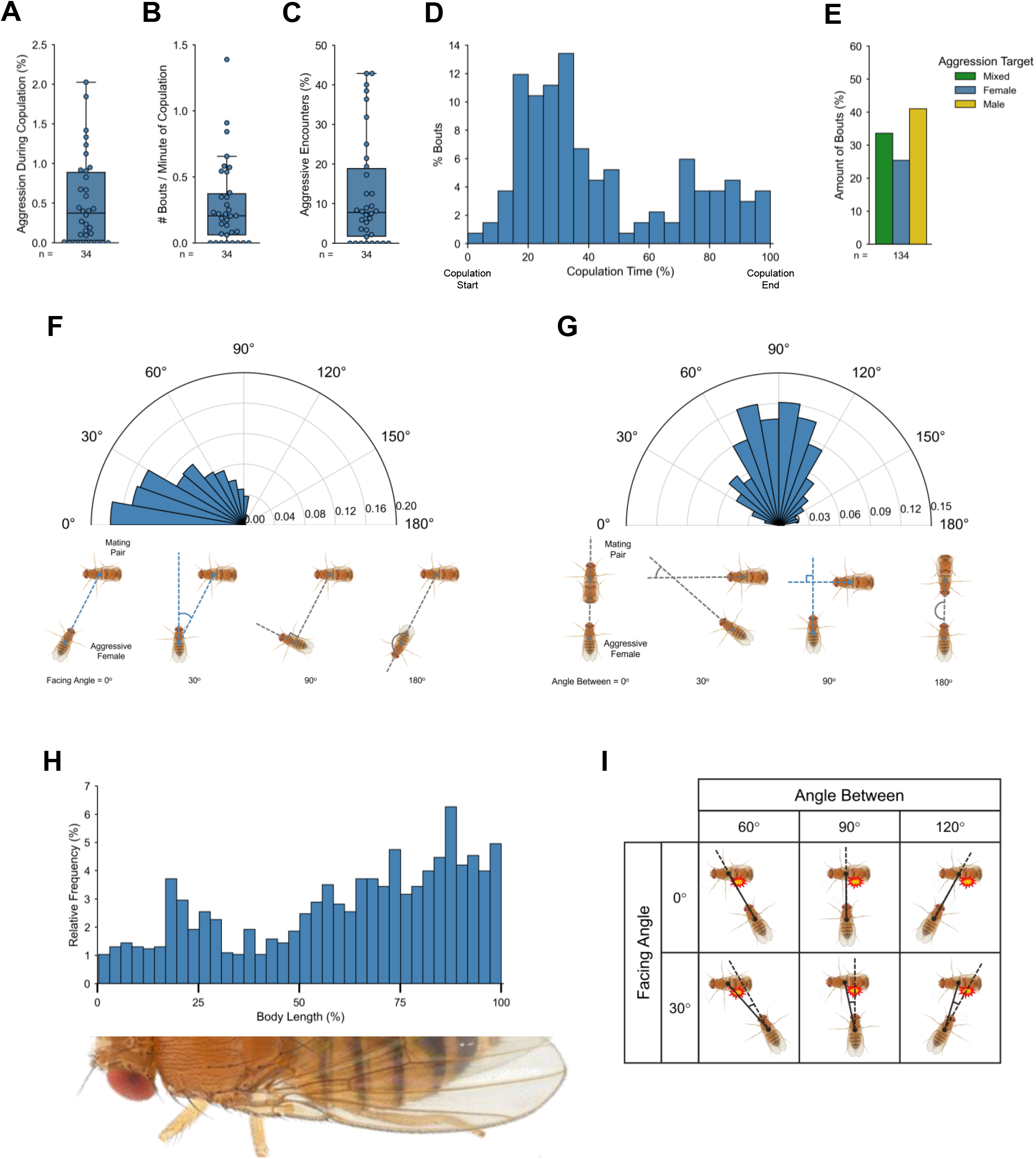
Female aggressive behaviour is reliable and stereotyped in the presence of a mating pair. **(A)** Aggression rate of wild type females towards a mating pair. **(B)** Number of aggression bouts per minute of copulation of wild type females towards a mating pair. **(C)** Percentage of encounters where aggression occurs. **(D)** Distribution of aggression occurrence during copulation. Copulation time is expressed as a percentage to normalize for different copulation durations. 0% represents the start of copulation and 100% the end of copulation, irrespective of copulation duration. n = 134 aggression bouts from 34 tested females. **(E)** Percentage of aggression bouts targeting the mating female, the mating male, or both. n = 134 aggression bouts from 34 tested females. **(F)** Distribution of facing angles. Top: the polar axis represents the range of possible angles, while the radial axis represents the percentage of total angles that fall within any given range. Angles are binned in 10-degree intervals. Bottom: schematic of representative facing angles. Blue lines match the range of angles found in the distribution. **(G)** Distribution of angles between flies. Top: the polar axis represents the range of possible angles, while the radial axis represents the percentage of total angles that fall within any given range. Angles are binned in 10-degree intervals. Bottom: schematic of representative facing angles. Blue lines match the range of angles found in the distribution. **(H)** Distribution of points along the mating female’s body axis that are targeted during aggression. The body axis is represented as the percentage of fly length to normalize for variation in female body size. **(I)** Schematic summarizing the information from facing angles (**H**), angles between (**I**), and body length hit position (**J**). The red-bordered yellow star represents the body area with the most hits. Sample size (number of flies, unless otherwise stated) is shown under each graph.

### Aggression does not affect copulation duration, latency to the second copulation, number of eggs laid, or hatching rate

From our previous findings (Figures 1D and 1H), we asked whether the observed aggressive displays could have some effect on the reproductive outcomes of either the target or aggressor female. To address this question, we compared three different contexts: mating pairs together with an aggressor female, such that aggressive interactions can occur (“with aggression”, see Movie S2); mating pairs separated from a competing female by a mesh, such that they share the same arena but aggressive interactions will not occur (“partition”, see Movie S3), the mesh being removed after copulation ended to allow the competitor female to also mate; or mating pairs by themselves, such that there is no aggression and mating pairs are not disturbed by any other potential signals coming from the competing female (“single couple”; see Star Methods). If the purpose of aggressive displays is to cause the male to dismount earlier from the current mating, possibly curtailing ejaculation, we would expect to see a reduction in the duration of the first copulation. This is not the case, as we found that copulation duration remains undisturbed in the presence or absence of aggression (Figure 2A). Alternatively, if aggressive displays are a strategy to prime the male so that the second copulation can start sooner, then we would expect to see a decrease in the latency to the second mating. However, we found that the latency to the second copulation is unaffected by the presence or absence of female aggression (Figure 2B). So far, we looked at potential short-term consequences of aggression. However, we reasoned that aggressive displays might have more long-term implications, specifically at the level of the number of eggs laid and progeny fitness. Aggressive behaviour could be used as a mechanism to lower the fitness of the competitor; additionally, such aggressive displays could incur a high metabolic cost and therefore directly impact the reproductive capacity of the aggressor. To explore this, we collected both females (target and aggressor) after each experiment and counted the number of eggs laid by each after 24 hours. In addition, we also counted the number of total adults ecloded from those laid eggs and used that to calculate the hatching rate (see Star Methods). Surprisingly, we found no difference either in the number of eggs laid, or their hatching rate of the target female in the presence or absence of aggression (Figures 2C and 2D), suggesting that being subjected to aggressive displays has no bearing on egg production or viability. Similarly, the number of eggs laid and their hatching rate of the potentially aggressor female were also unaffected in the presence or absence of aggression (Figures 2E and 2F), implying that even if aggression is energetically taxing, it is not to the point of limiting egg production or viability. Although we could not uncover the biological consequences of this aggressive behaviour, the consistency with which we can observe it and the stereotyped nature of its execution hint at its biological relevance.

**Figure 2.**
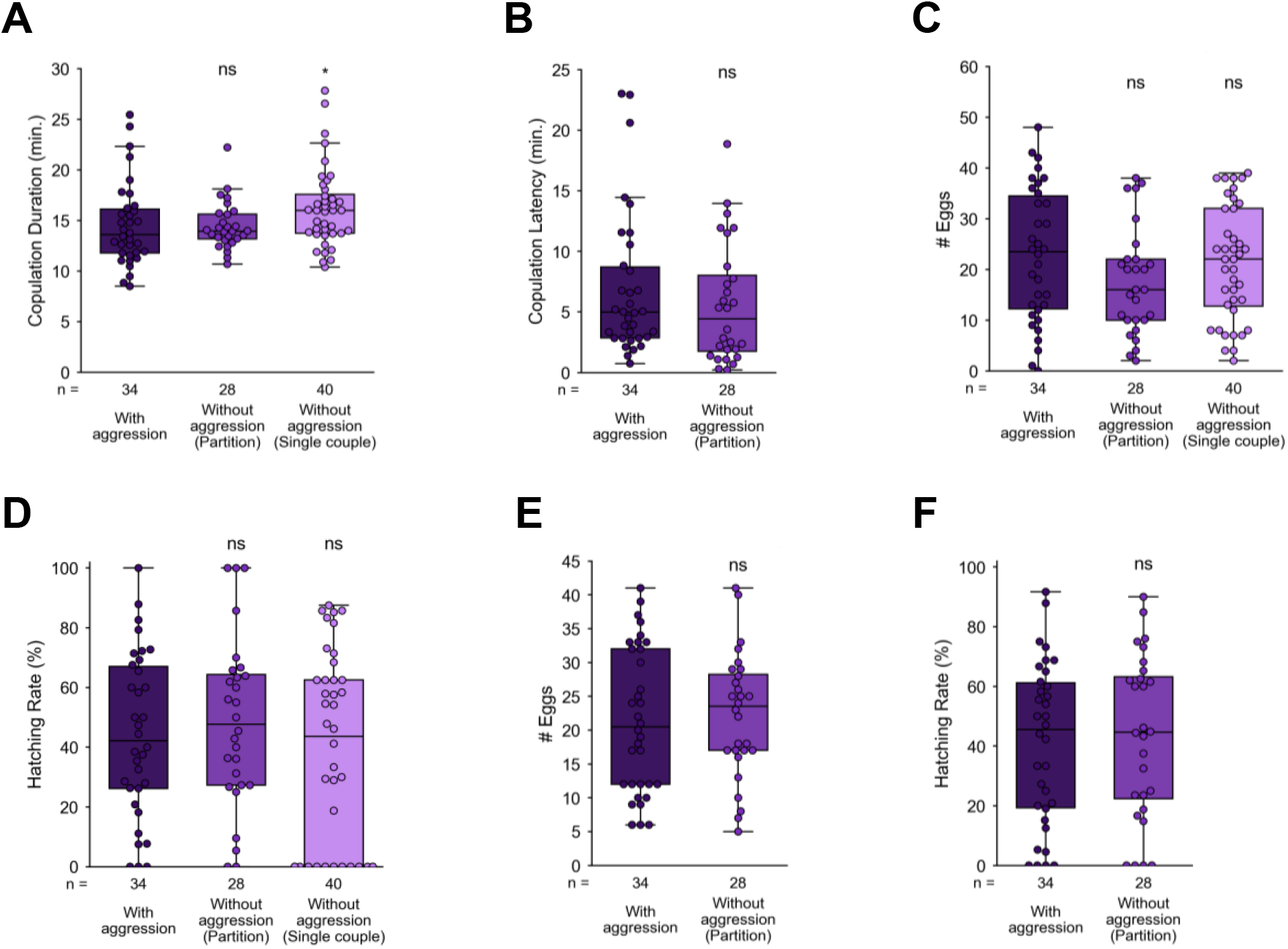
Aggression does not affect copulation duration, latency to the second copulation, number of eggs laid, or hatching rate. **(A)** Duration of the first copulation (target female). **(B)** Latency to the second copulation (competitor female). **(C)** Number of eggs laid from the first copulation (target female). **(D)** Hatching rate from the first copulation (target female). **(E)** Number of eggs laid from the second copulation (competitor female). **(F)** Hatching rate from the second copulation (competitor female). See Supplementary Table 1 for p-values and effect sizes; ns = not significant, *p < 0.05. Sample size (number of flies tested) is shown for each condition below its corresponding boxplot.

### Mating drive regulates aggressive behaviour

The fact that females are aggressive during the first copulation, but no aggression is observed by the recently mated female towards the second mating pair lead us to hypothesize that female aggressive drive might be modulated by the mating status of the female, one component of the animal’s internal landscape. The behavioural changes associated with the transfer of sex peptide during mating are well documented, leading, for instance, to drastic shifts within 24 hours in egg laying, receptivity, feeding preference, and even fighting for food patches^31,41–45^. It is therefore reasonable that the same, or similar mechanisms might be at play here, modulating aggressive behaviour between the virgin and mated states of the female. Additionally, it has been recently reported that the experience of copulation itself is enough to lead to early onset of behavioural changes, although through an independent mechanism than that of the sex peptide pathway^46^. To test whether the mating status changes affect female aggressive displays, we quantified the aggression rate of 24h-mated females, as well as 2h-mated females, respectively. We found that in both cases female aggression levels were strongly reduced (Figures 3A, S2A, and S2B). This effect is not simply due to a general lack of activity of mated females, since these animals walk the same distance as virgin controls (Figure 3B), suggesting that not only the sex peptide-mediated changes in female physiology, but also mating experience are sufficient to significantly and specifically impact aggressive drive. Is the post-mating reduction in female receptivity, and presumably in the mating drive, the leading cause for lower aggression? To test this, we decided to study the effect of courtship deprivation on female aggressive drive. We quantified the aggression rate of virgin females that were not courted by a male (i.e., courtship deprived) and that were only introduced in the behavioural arena during the copulation of a male-female pair. We observed that depriving the females of courtship is also enough to significantly reduce aggressive displays (Figures 3A, S2A, and S2B). These results show that mating drive and aggression drive are positively associated in females.

**Figure 3.**
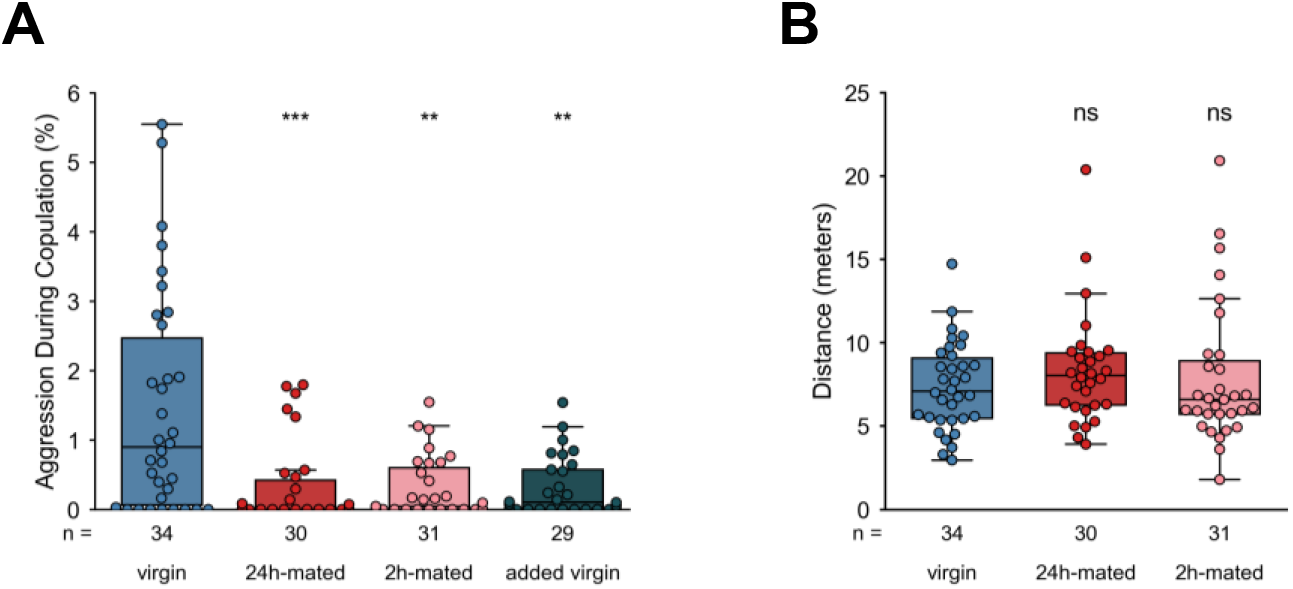
Mating drive regulates aggressive behaviour. **(A)** Aggression rate of wild type virgin, 24h-mated, 2h-mated, and added virgin females towards a mating pair. **(B)** Distance walked by non-mating females, from the start of the experiment until the end of copulation. See Supplementary Table 1 for p-values and effect sizes; ns = not significant, **p < 0.01, ***p < 0.001. Sample size (number of flies tested) is shown for each condition below its corresponding boxplot.

### Olfaction is required for normal levels of aggression

Having pinpointed one component of the internal state of the female fly that regulates its aggressive behaviour, we next sought to investigate which elements in the external environment of the fly could influence this behaviour. To address this question, we tested the contribution of gustation, hearing, vision, and olfaction with the use of either mutant or otherwise manipulated flies (see Star Methods). We found that visual mutants and, more strikingly, olfactory mutants show a decrease in aggressive behaviour compared to wild type females, stressing the importance of these two modalities (Figures 4A, S3A, and S3B). We have shown before that aggressive displays are a targeted behaviour, i.e., females orient themselves towards their intended target (Figure 1I). However, if that is the only role played by vision in aggression, then we would expect to see females displaying aggression when they are not in proximity of the mating pair. We found that blind females display aggression exclusively towards the mating pair, suggesting that vision is likely gating aggression in some other way, perhaps allowing females to visually recognize a mating pair of flies. The effect of removing hearing did not reach statistical significance, but we do note that it is nonetheless quite large, reducing aggressive displays by around 75% (Figures 4A, S3A, and S3B; see Supplementary Table 1). This is in line with previous reports of hearing regulating aggressive behaviour in fruit flies^21^. One possibility that explains this effect is that acoustic stimulation by courtship song increases the female mating drive, and by removing hearing, and therefore keeping mating drive low, aggression drive plummets accordingly. Alternatively, aggressive displays may be partially driven by the detection of the mating female’s song, given that recent studies have shown that females sing during copulation^47^. Only the removal of gustation clearly shows no effect on female aggression (Figures 4A, S3A, and S3B). We also found that the aggression effects observed in each of the sensory conditions are not a reflection of the flies’ general lack of activity, since none of the conditions walks a significantly different distance than wild type control flies (Figure 4B). Therefore, olfaction seems to be the primary sensory modality that females require in order to identify the appropriate environment in which to perform aggressive displays.

**Figure 4.**
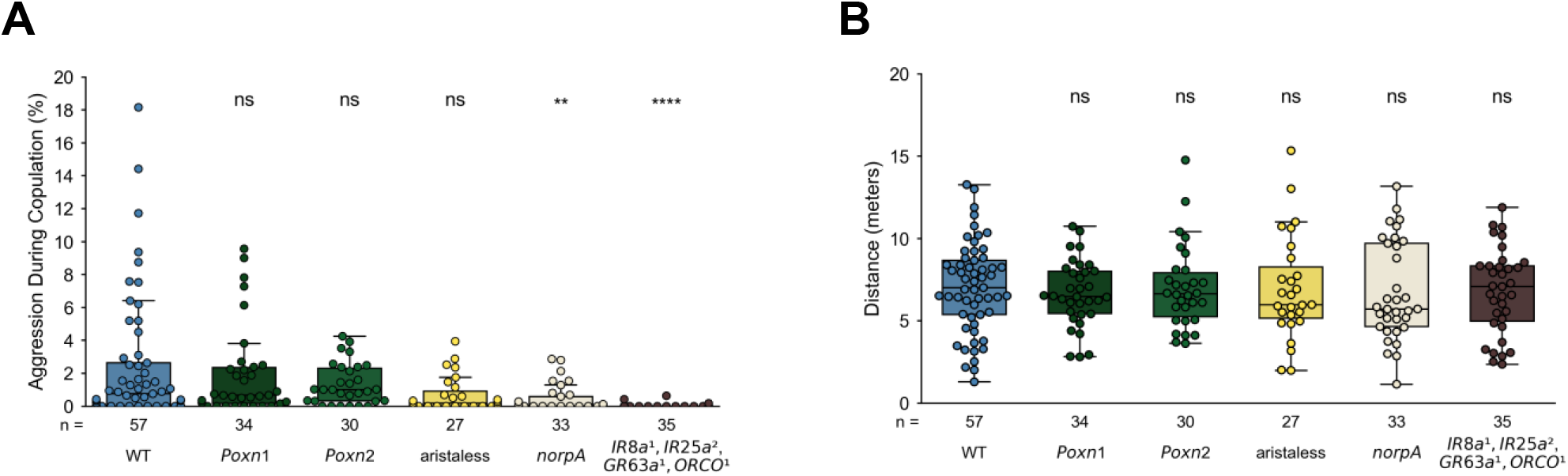
Olfaction is required for normal levels of aggression. **(A)** Aggression rate of wild type, tasteless, deaf, blind, and anosmic virgin females towards a mating pair. **(B)** Distance walked by non-mating females, from the start of the experiment until the end of copulation. See Star Methods for fly genotypes, and Supplementary Table 1 for p-values and effect sizes; ns = not significant, **p < 0.01, ****p < 0.0001. Sample size (number of flies tested) is shown for each condition below its corresponding boxplot.

### Presence of food odour and activity of OR47b olfactory sensory neurons contribute to female aggressive behaviour

We next sought to narrow down the number of possible candidates within the olfactory landscape of the female fly that may modulate aggressive behaviour. Customarily we lace all arenas with food paste prior to experiments, which is then removed, leaving only an odour trace of the food in the arenas (see Star Methods). Food odour has been shown to stimulate male courtship^48^, an important part of our experiments, which depend on successful copulation. Since males are stimulated by the smell of food and given that both males and females fight for territories^3,21,29,31,39^, we reasoned that food odour could be responsible for regulating female aggressive behaviour. To test this hypothesis, we compared the aggression rate of females in the presence of a mating pair, either with or without food odour. We found that food odour is indeed required for aggressive behaviour to be displayed (Figures 5A and S4A). This could mean that food odour acts as a proxy for the presence of a territory and, therefore, that the behaviour we have characterized thus far is in fact a fight for territory. If this is the case, then removing the male, and therefore copulation, should yield high levels of aggressive behaviour in the presence of food odour only. We found that in the absence of a mating pair, exposure to food odour by itself is not enough to induce the high levels of aggression observed in the presence of both components (compare Figures 5B and S4B with Figures 5A and S4a). This suggests that food odour together with the presence of a mating pair are needed for high number of aggressive displays to take place. This implies that some additional olfactory cue from the mating pair is needed in addition to that of food odour. Olfactory and ionotropic receptors have extensively been shown to regulate many fly social behaviours, including reproductive and aggressive behaviours^45,48–55^. However, the fly olfactory system is comprised of dozens of such receptors. Hence, we needed an approach to narrow down the number of potential candidates. It is well established that the *Drosophila* fruitless gene encodes sexually dimorphic behaviour by specifying sexually dimorphic circuits during development^22,56,57^. This dimorphism is reflected in many behaviours, one of which being aggression^9,18,25^. Three *Drosophila* receptors express *fruitless*^58,59^: IR84a, which senses a specific range of chemicals present in the odour blend of food, and shown to be responsible for the food-stimulating effect on male courtship^48^; OR67d, which detects the male-specific pheromone 11-cis-Vaccenyl Acetate (cVA), shown to be a strong mediator of male-male aggression^23,51^; and OR47b, which senses compounds present in both males and females, and is a known modulator of receptivity in females and courtship in males^49,50,52,54,55,60^. When we silenced the sensory neurons expressing each of the individual receptors, we found that only OR47b showed a significant decrease of around 40% in aggression rate (Figures 5E and S4E). Silencing IR84a had no effect on female aggression (Figures 5C and S4C). Given that IR84a is tuned to detect a very narrow range of food-related odours, these findings indicate that other olfactory receptors are responsible for the food odour-derived modulation of aggression. Finally, silencing OR67d also did not affect aggression rate in a significant way (Figures 5D and S4D), suggesting that, surprisingly, the male-specific olfactory cue, cVA, seems to not be necessary for the proper display of female aggressive behaviour. We conclude from these results that multiple olfactory cues are required for the proper expression of female aggressive behaviour, specifically, those from food, via as-yet uncharacterized receptors, and those form other flies, through the activity of OR47b-expressing olfactory neurons.

**Figure 5.**
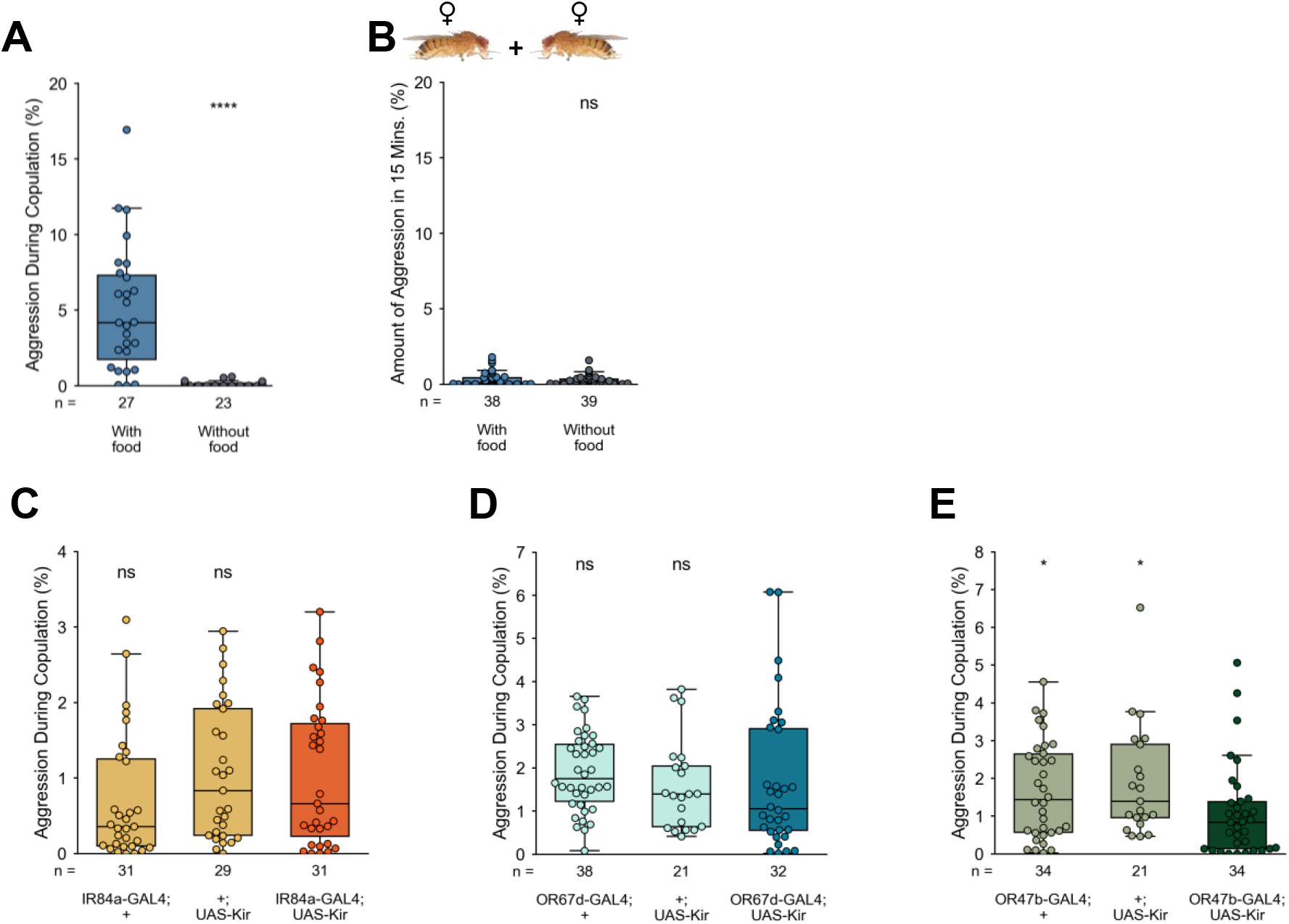
Presence of food odour and activity of OR47b olfactory sensory neurons contribute to female aggressive behaviour. Aggression rate in the following conditions: **(A)** Wild type flies in the absence or presence of food odour. **(B)** Wild type female pairs, without a male, in the absence or presence of food odour. **(C)** Silenced IR84a flies and respective controls in the presence of food odour. **(D)** Silenced OR67d flies and respective controls in the presence of food odour. **(E)** Silenced OR47b flies and respective controls in the presence of food odour. See Star Methods for fly genotypes, and Supplementary Table 1 for p-values and effect sizes; ns = not significant, *p < 0.05, ****p < 0.0001. Sample size (number of flies tested) is shown for each condition below its corresponding boxplot.

## DISCUSSION

Aggressive behaviour occurring in the context of intrasexual competition is an important trait for animal fitness as it allows animals to compete for limited resources. In this study, we have found that females of *Drosophila melanogaster* will compete for mates, specifically by engaging in aggressive displays towards mating pairs. These displays are stereotyped and strongly dependent on olfactory cues from both food and other flies, with a significant, yet partial, contribution of OR47b sensory neurons.

Previous work in female *Drosophila* aggression has focused on competition over territory, that is, where two females are fighting over a physical patch of food. Interestingly, in that context mated females are more aggressive than virgin females^31,32,37^, in stark opposition with our own findings in the context of mate competition. This reversal in the relationship between mating status and aggression levels could be due to a shift in the females’ perceived value of available resources: virgin females would value mating partners higher than egg-laying sites, whereas mated females would then prioritize egg-laying site acquisition over possible early rematings. Indeed, it has been well established that the mating status of femal*e Drosophila* is capable of reversing the valence of resources, with mated females displaying a strong dietary yeast and salt preference in contrast to the significant sugar preference of virgin counterparts^43,44^. It is therefore reasonable that a similar mechanism might regulate female aggression levels as a function of mating status and available resources.

Our finding that food odour, in addition to the presence of a mating pair, is required to drive aggression in virgin females suggests that this behaviour is contingent on an ecologically relevant context, as either cue by itself fails to elicit appropriate aggressive responses. Whether this is because, as reported in males^48^, food odour has a stimulatory effect that increases female mating drive, or the odour itself is representative of a nearby egg-laying site remains to be elucidated.

Our study reveals a strong dependence of aggressive behaviour on olfaction, largely contributed to by OR47b, which is tuned to a select few elements of flies’ cuticular hydrocarbon profiles. A growing body of work has been uncovering the many roles this olfactory receptor plays in mediating *Drosophila* reproductive interactions, from young female preference and mating advantage in males, to increased mate choosiness in females^50,52,55,61^. In fact, the behavioural differences we observed between virgin and mated females fit very well with the described physiological modulation of OR47b OSNs occurring after mating. Briefly, juvenile hormone production induced by mating alters the sensitivity of OR47b neurons, changing the behavioural output to the same sensory input depending on the age or mating status of the individual. Just as desensitized OR47b neurons impart increased choosiness in females, this mechanism could just as easily gate aggressive behaviour by dampening aggression-triggering olfactory stimuli in mated females. Virgin females, on the other hand, would integrate the information provided by more sensitive OR47b neurons with their internal mating drive in order to express an appropriate aggressive response.

While using anosmic females lead to a complete abrogation of aggressive behaviour, removing OR47b produced only a partial reduction in female aggression. It is clear that additional olfactory sensors are involved in evaluating the correct sensory context in which to execute aggressive behaviour. Given our findings that food odour is essential for driving aggression, these additional receptors are most likely ones tuned to relevant food odour components. Besides olfaction, vision and, likely, hearing were also revealed to have an impact in female aggression levels. This is in line with previous work reporting the involvement of these modalities in male aggression^14,16,21^, suggesting even more parallels with female aggression. Males require a moving object to express aggression^14^, whereas in females vision is likely used to distinguish the general shape of mating rivals, since we failed to observe any instance of aggression targeted anywhere but towards the mating pair. As for acoustic signals, it has been reported that males respond with aggressive displays to agonistic sounds of rival males^21^. In the context of female mate competition, aggressive females might be responding to the copulation song generated by the mating female^47^. Thus, females might be using acoustic, visual, and olfactory cues as long-, mid-, and close-range signals to identify potential competitors and execute appropriate aggressive levels towards them. Whether such a system is actually being employed by females, and which specific features of the environment each modality is picking up to contribute to female aggressive displays remains unclear.

pC1 is a known key integrator node that regulates reproductive and aggressive behaviours in *Drosophila*, with different cell types contributing to different behaviours^11,12,20,62–64^. pC1d in particular has been shown to be responsible for driving aggressive displays in females, with other cell types mediating functions related to receptivity^12^. In addition to this, recurrent activity of pC1 has been shown to generate a persistent internal state in the fly brain, thus providing a neural basis for context-specific stimulus integration. It is therefore likely that this same circuitry is being recruited in the context of our study, regulating mate-driven aggressive competition and possibly weighing the contribution of olfactory, auditory, and visual sensory inputs together with the internal state of the female.

Many fruit flies exhibit lekking mating systems, including Tephritidae and Hawaiian *Drosophila* species. These mating systems are characterized by four main features: 1) males provide no parental care, and supply only gametes; 2) males are spatially aggregated in mating areas, or leks; 3) males do not control access to resources critical to females; and 4) females are free to select mates at the lek^65^. Although field evidence is lacking regarding whether *Drosophila melanogaster* is a true lekking species, it does exhibit some of its elements. Despite being uncommon, this type of mating system is taxonomically widespread, being present in other insects, crustaceans, fish, reptiles, birds, and mammals^33,35,36,65–72^. In these systems, males will perform courtship displays to females, who visit the leks for the sole purpose of mating. Under these circumstances, interfemale competition, many times in the form of aggression, takes place. However, the biological significance of these aggressive behaviours is still poorly understood, although several hypotheses have been put forth. One reason for aggression may be to reduce the waiting time for access to a preferred male, where the cost of aggression would be lower than delaying mating with a high fitness partner or mating with a lower fitness partner. Alternatively, aggression could be used as a way to reduce the fitness of rival females by, for example, disturbing copulation, in a classic case of spiteful behaviour. Finally, aggressive females might compete for high quality or quantity of sperm in a limited environment, therefore attempting to decrease sperm volume transferred to competitors. In our work we report no effect of female aggression on copulation duration, nor on the reproductive output of either female involved. It therefore seems that none of the strategies offered above are at play here. Indeed, active copulation disruption induced by female aggression seems rare across species^33,35,36^. Even in the case of the Mediterranean fruit fly, *Ceratitis capitata*, where virgin females display intense aggression towards mating pairs, the advantage of executing such displays remains elusive.

In conclusion, we report here how female aggression can be elicited by mating pairs in the presence of food odour and highlight the importance of social context in the characterization of behaviour. These findings pave the way for addressing the neural underpinnings of female aggressive behaviour and add to the growing body of evidence that *Drosophila* females display rich, complex behaviours, that are sensitive to social, environmental, and internal state.

## Supporting information

Key Resources Table

## ACKNOWLEDGEMENTS

We thank João Frazão for help with Bonsai software, Hugo Cachitas and Alexandre Laborde for help with Python analysis, Cecilia Mezzera from the Vasconcelos lab for help with manual annotations, Alexandre Azinheira for help with video editing and preparation, Kathrin Steck from the Ribeiro lab for sharing the chemosensory mutant lines, and Eyrún Eyjólfsdóttir from the Computer Vision lab at Caltech for help with FlyTracker. We are also very thankful to the many platforms at Champalimaud Foundation: the Fly Facility for dissections, immunostainings, and fly food; the Scientific Software Platform for developing and helping with PythonVideoAnnotator; the Scientific Hardware Platform for support with and developing custom electronic HARP behaviour boards. Finally, we thank Cristina Ferreira from the Vasconcelos lab for feedback on the manuscript. This work was supported by Fundação Champalimaud, by Portuguese national funds, through FCT - Fundação para a Ciência e a Tecnologia - in the context of the project UIDB/04443/2020, a grant awarded to M. G. - PD/BD/105943/2014 -, the research infrastructure Congento, LISBOA-01-0145-FEDER-02270, and BioData.pt - LISBOA-01-0145-FEDER-022231. *Drosophila melanogaster* images used throughout this work were taken from Nicolas Gompel’s lab webpage (http://gompel.org/images-2/drosophilidae) under a Creative Commons license.

## AUTHOR CONTRIBUTIONS

Conceptualization, M. G. and M. L. V.; Methodology, M. G. and M. L. V.; Software, M. G.; Formal Analysis, M. G.; Investigation, M. G.; Resources, M. G., S. D. and M. L. V.; Data Curation, M. G. and S. D.; Writing – Original Draft, M. G. and M. L. V.; Writing – Review & Editing, M. G. and M. L. V.; Visualization, M. G.; Supervision, M. L. V.; Project Administration, M. L. V.; Funding Acquisition, M. G. and M. L. V.

## STAR⋆METHODS

## RESOURCE AVAILABILITY

### Lead contact

Further information and requests for resources should be directed to and will be fulfilled by the Lead Contact, Maria Luísa Vasconcelos.

### Materials availability

This study did not generate new unique reagents. The materials built in-house are open-source and can be made available through our institute’s scientific hardware platform (Champalimaud Hardware Platform; http://www.cf-hw.org/).

### Data and code availability

Raw movies acquired in the current study have not been deposited in a public repository because of their large size, but annotation files in csv format generated from acquired movies, as well as tracking output files and necessary Python code used to generate all results present in the current study have been deposited on Harvard Dataverse and are publicly available as of the date of publication of this manuscript. DOIs are listed in the key resources table. Original, raw movies, as well as additional tracking output files or further information required to interpret the data reported in this paper are available on request from the corresponding author.

## EXPERIMENTAL MODEL AND SUBJECT DETAILS

Fruit flies *D. melanogaster* were raised in standard cornmeal-agar medium, using Vienna food recipe (in 1L of water: 80g molasses-barley malt, 22g beet syrup, 80g corn flour, 18g granulated yeast, 10g soy flour, 8g agar-agar, 8mL propionic acid, 12mL 15% nipagin, 35mL Bavistin), at 25ºC and 70% relative humidity in a 12h dark:12h light cycle. For information on fly stocks used, please refer to the Key Resources Table; for detailed information on fly genotype for each experiment, please see Star Methods.

## METHOD DETAILS

### Fly genotypes

The detailed genotypes per figure are as follows:

Figure 1:

Wild type (DL strain)

Figure 2:

Wild type (DL strain)

Figure 3:

Wild type (DL strain)

Figure 4:

Control: wild type (DL strain)

Poxn1 mutant: *w*^*1118*^ ; *PoxnΔ*^*[M22-B5]*^ ; *Δ*^*[SfoBs105]*^ */ Δ*^*[SfoBs127]*^

Poxn2 mutant: *w*^*1118*^ ; *PoxnΔ*^*[M22-B5]*^ ; *Δ*^*[SfoBs127]*^ */ Δ*^*[SfoBs105]*^

Aristaless: wild type (DL strain) with both aristae removed

Blind mutant: *w[*], norpA*^*36*^ ; *+* ; *+*

Anosmic mutant: *IR8a*^*1*^; *IR25a*^*2*^; *Orco*^*1*^, *GR63a*^*1*^

Figure 5a-b:

Wild type (DL strain)

Figure 5c:

Control 1: *w[*]* ; *IR84a-GAL4 / +* ; *+*

Control 2: *w[*]* ; *+ / UAS-Kir2*.*1-EGFP* ; *+*

Test: *w[*]* ; *IR84a-GAL4 / UAS-Kir2*.*1-EGFP* ; *+*

Figure 5d:

Control 1: *w[*], OR67d-GAL4 / +* ; *+* ; *+*

Control 2: *w[*]* ; *+ / UAS-Kir2*.*1-EGFP* ; *+*

Test: *w[*], OR67d-GAL4 / +* ; *+ / UAS-Kir2*.*1-EGFP* ; *+*

Figure 5e:

Control 1: *w[*]* ; *OR47b-GAL4 / +* ; *+*

Control 2: *w[*]* ; *+ / UAS-Kir2*.*1-EGFP* ; *+*

Test: *w[*]* ; *OR47b-GAL4 / UAS-Kir2*.*1-EGFP* ; *+*

Figure S1:

Wild type (DL strain)

Figure S2:

Wild type (DL strain)

Figure S3:

Control: wild type (DL strain)

Poxn1 mutant: *w*^*1118*^ ; *PoxnΔ*^*[M22-B5]*^ ; *Δ*^*[SfoBs105]*^ */ Δ*^*[SfoBs127]*^

Poxn2 mutant: *w*^*1118*^ ; *PoxnΔ*^*[M22-B5]*^ ; *Δ*^*[SfoBs127]*^ */ Δ*^*[SfoBs105]*^

Aristaless: wild type (DL strain) with both aristae removed

Blind mutant: *w[*], norpA*^*36*^ ; *+* ; *+*

Anosmic mutant: *IR8a*^*1*^; *IR25a*^*2*^; *Orco*^*1*^, *GR63a*^*1*^

Figure S4a-b:

Wild type (DL strain)

Figure S4c:

Control 1: *w[*]* ; *IR84a-GAL4 / +* ; *+*

Control 2: *w[*]* ; *+ / UAS-Kir2*.*1-EGFP* ; *+*

Test: *w[*]* ; *IR84a-GAL4 / UAS-Kir2*.*1-EGFP* ; *+*

Figure S4d:

Control 1: *w[*], OR67d-GAL4 / +* ; *+* ; *+*

Control 2: *w[*]* ; *+ / UAS-Kir2*.*1-EGFP* ; *+*

Test: *w[*], OR67d-GAL4 / +* ; *+ / UAS-Kir2*.*1-EGFP* ; *+*

Figure S4e:

Control 1: *w[*]* ; *OR47b-GAL4 / +* ; *+*

Control 2: *w[*]* ; *+ / UAS-Kir2*.*1-EGFP* ; *+*

Test: *w[*]* ; *OR47b-GAL4 / UAS-Kir2*.*1-EGFP* ; *+*

### Behavioural experiments

For all experiments both male and female flies were collected under CO_2_ anaesthesia and raised in isolation at 25ºC and 70% relative humidity and aged 4-8 days until the day of the experiment. All experiments were performed at 25ºC, 70% relative humidity, in dim light, and between Zeitgeber times 0 and 4. For Figures 1, 3, 4, S1, S2, and S3 experiments, all flies were collected as late-stage pupae. For practical purposes, all flies were collected as early adults (1-3 hours post-eclosion) for all other experiments, since we found no difference in behaviour between the two collection methods (data not shown). All flies were inspected both at collection time and briefly before the start of experiments for any noticeable physical defects. Any fly that exhibited any of the following traits were discarded from being used in experiments: broken or bent tarsi, malformed legs, damaged or curled wings, wings locked at non-resting positions, bloated abdomens, abnormal walking patterns and markedly reduced walking. To distinguish between the two females, we selected one of the females in each pair at random and painted them over the posterior half of the thorax and the scutellum with a metallic silver 0.8mm nib roller-ball pen (Uni-Ball Signo UM-120NM). To hold the females in place during painting, they were pinned by one of their midlegs using precision forceps (Fine Science Tools Dumont #5CO, item nº11295-20), applying as little force as necessary to avoid damage to the legs. To limit the effect of this procedure as a possible confounding factor, we also subjected males and unpainted females to CO_2_ anaesthesia and manipulated them with the same precision forceps. After the painting procedure, all flies were allowed to recover at 25ºC and 70% relative humidity for at least 36 hours before experiments. Unless stated otherwise, all experimental arenas were laced with fly food paste at least overnight to imbue the arenas with the smell of food to stimulate courtship. To prepare this paste, we added 1mL of milliQ water to a regular fly food vial of standard cornmeal-agar medium and physically mashed the food and water together until a consistent paste was formed. A small amount of this paste was transferred to each conical arena, or enough was transferred to fill the smaller, rectangular arenas. This paste was removed with the aid of paper towels as thoroughly as possible prior to aspirating flies into the arenas to start the acquisition of experiment movies.

Figure 1 and S1 experimental data originates from the same experimental group and dataset as that of Figures 3 and S2.

For Figure 2 experiments, to ensure a lack of any aggressive behaviour towards the mating pair, we introduced a partition in the arenas (see Behavioural arenas section below) to physically separate the mating pair from the non-mating female. Briefly, the experiments start with partition in place, and one male with one female are gently aspirated into one of the partition sides, while the second female is aspirated into the other side of the partition. After 30 minutes, which ensures mating has occurred, the partition is removed to allow the isolated female to mate. At the end of the experiment, each of the two females in each arena was gently aspirated from the arenas at the end of the experiments and transferred to a vial of standard cornmeal-agar medium (using Vienna food recipe) and kept at 25ºC and 70% relative humidity. After 24 hours, if the females were still alive, they were discarded, the number of eggs laid in the vial was counted, and the vials were returned to 25ºC for another 9 days. At that point we counted the number of ecloded adults, keeping the vials at 25ºC and counting newly ecloded adults every day for 4 to 6 days additional days. To ensure that both females were mated during the experimental period, movies were acquired for 1 hour.

For Figure 3 and S2 experiments, to generate 24h-mated females we added a naïve male to the females’ vials 24 hours before the start of next day’s experiments. To ensure that flies were mated, the females’ vials were kept and incubated at 25ºC and 70% relative humidity for 10 days and checked for ecloded progeny. To generate 2h-mated females, naïve males were added to the females’ vials at the start of experiments, and left undisturbed for 2 hours, after which they were included in the last batch of experiments of the day. Mating occurrence was checked visually. For the added virgin condition, for each experiment we started by gently introducing a single virgin female and a single naïve male to the arena, then checked visually for the start of copulation, upon which we then gently aspirated a second virgin female into the same arena where the initial pair just started mating. Movies were acquired for 1 hour.

For Figure 4 and S3 experiments, to remove gustation, we crossed two different homozygous Poxneuro genetic deletions to each other in two both directions (PoxN1 and PoxN2); to remove vision we used a homozygous mutant for the norpA gene; to remove olfaction, we employed a quadruple mutant for both of the ionotropic receptor co-receptors, IR8a and IR25a, the olfactory receptor co-receptor, OR83b, and the CO_2_ receptor GR63a (see Star Methods for detailed fly genotypes); finally, to remove hearing, we removed both aristae in female flies. To do this, individual flies were anesthetized with CO_2_ approximately 24 hours before the experiment. Aristae were cut bilaterally at their base with micro scissors (World Precision Instruments) under a scope. Flies were allowed to recover at 25ºC and 70% relative humidity until the experiment. Movies were acquired for 45 minutes.

For Figures 5a, 5b, S4a, and S4b experiments, the “food” conditions are performed as all previous experiments, i. e., by lacing arenas with food paste prior to testing flies. For the “no food” conditions, arenas are not laced with the food paste. In 5b and SS4b, since there is no copulation from which to take the first 5 minutes for analyses, we instead annotated aggression during the first 15 minutes of the experiment. Movies were acquired for 55 minutes. For Figures 5c, 5d, 5e, S4c, S4d, and S4 experiments, IR84-, OR67d-, and OR47b-expressing olfactory sensory neurons were silenced using the inwardly rectifying potassium channel Kir2.1 that hyperpolarizes the neurons, thus preventing action potential formation^73^. Movies were acquired for 30 minutes.

### Behavioural arenas

Figures 1, 3, 4, S1, S2, and S3 experiments were recorded in a 2×2 array of custom-made circular arenas with a conical-shaped bottom, as previously described^74,75^. These arenas are made of mechanically bored white polyoxymethylene, with 11º sloped walls, 4mm maximum height, and approximately 3cm of walking diameter, topped with clear acrylic lids. Figures 2, 5 and S4 experiments were recorded in a 4×4 array of 20×17mm rectangular, clear acrylic arenas with approximately 17×10mm of walking area and 3mm height, topped with clear acrylic lids. For Figure 2 experiments, we constructed additional rectangular arenas with the same specifications as before but leaving a 0.6mm-wide slit on both sides of the arenas. Through these openings a nylon plastic mesh (SEFAR-NITEX® 06-500/38) was placed, separating one of the females from the other female and male. This partition can then easily be removed at any time during movie acquisition without interrupting the experiments.

## QUANTIFICATION AND STATISTICAL ANALYSIS

### Movie acquisition

Experiments were recorded at 60 frames per second with a camera (Point Grey Flea®3 FL3-U3-32S2M) equipped with a 5mm MegaPixel fixed focal length lens (EdmundOptics®, stock nº64-867) mounted above the arenas. Movies were acquired in dim light using 940nm LEDs integrated in a custom electronic LED array board (LED array v3.0) with associated control board (LED array interface v1.0) and control software (HARP version v0.3) designed and built in-house (Scientific Hardware Platform) and a UV/VIS cutoff M43.0×0.75 machine vision filter (EdmundOptics®, stock nº89-839). Flies were recorded in grayscale at 60 frames per second. Bonsai^76^ (version 2.4.0) was used to acquire the movies. Figures 1, 3, 4, S1, S2, and S3 experiments were recorded with a resolution of 960×940 pixels. Figures 2, 5 and S4 experiments were recorded with a resolution of 1248×1010 pixels.

### Data processing

After movies related to Figures 1, 3, 4, S1, S2, and S3 were acquired, FlyTracker^77^ was used to track the three flies and output information concerning their position, orientation, velocity, distance to the other fly, facing angle, and angle between flies. Because FlyTracker only provides angular and distance information for two animals, we adapted its code to ensure that these features were made available for our three animal assays (see Key Resources Table for code repository and respective DOIs). MATLAB version R2017a vas used to run FlyTracker. Movies related to Figures 2, 5, and S4 were tracked using the simpler algorithm from the in-house developed PythonVideoAnnotator (https://biodata.pt/python_video_annotator, version 3.306). Following tracking, PythonVideoAnnotator was used to manually annotate the time and duration of copulation and aggression bouts. Aggression bouts were annotated as any moment where continuous headbutt or shoving (the highest intensity aggressive displays in females^39^) were observed, in-between which wing flicking, and intense fencing could occasionally also be included. Bouts were considered separate instances when the flies would stop interacting for at least 1 second. Cases where the flies were in proximity but not interacting or only fencing were not classified as aggression.

### Quantification of behaviours

Data analysis was performed using custom Python 3.6.2 scripts for all experiments. For Figure 1 experiments, aggression during copulation was calculated as follows:

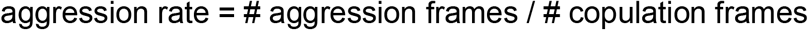

and the number of aggression bouts per minute as follows:

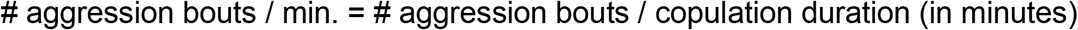

For all other experiments, we confirmed that the first 5 minutes of copulation are representative of the overall aggressive behaviour displayed during the entirety of copulation (data not shown). Therefore, all aggression metrics were calculated and analysed within the first 5 minutes of copulation. Aggression during copulation was calculated as follows:

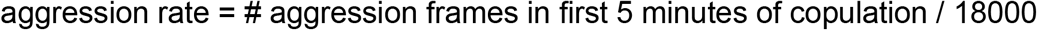

and the number of aggression bouts per minute as follows:

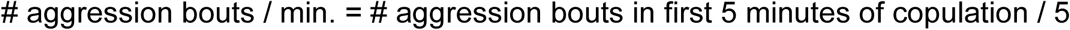

Experiments with aggression are calculated as the percentage of experiments where at least one bout of aggression occurred. Encounters are defined as moments where the non-mating female is within 4mm of the mating pair. Aggressive encounters are calculated as the percentage of encounters in which at least one aggression bout occurs. The number of aggressive bouts targeted towards males, females, or both was scored manually, and then converted to a percentage of the total amount. To get the body points along the mating female’s body targeted by aggressive females, we calculated the intersection of the aggressive female’s heading with the ellipsoid representation of the mating female’s body. This point was then converted to and angular representation to make it female body size-agnostic, and this angle can then be mapped to the mating female’s body axis. Distance walked was calculated from the start of each experiment until the end of the first copulation. We found no difference between the amount of distance walked before copulation starts and the amount of distance walked during copulation for the non-mated females (data not shown). The hatching rate is calculated as follows:

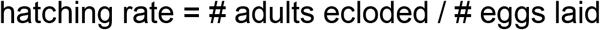

For all boxplots, the outline of the box represents the interquartile range (IQR), the upper whiskers are drawn up to Q3 + (1.5 x IQR), and the lower whiskers are drawn down to Q1 - (1.5 x IQR). The line inside the box denotes the median of each sample. Each dot on the overlapping swarm plot corresponds to a fly.

### Statistical analysis

Prior to statistical testing, any outliers for aggression rate were removed from our samples. Individual data points were considered outliers if, and only if, they both lied outside the whisker range of the boxplots and their absolute z-score was equal to or higher than 3 standard deviations. Outliers never comprised more than 6% of our total samples and were excluded from any all analyses. After discarding outliers, Levene’s test was used to assess variance homogeneity, and Shapiro-Wilk and D’Agostino-Pearson tests were used to assess normality across all individual experiments. For all pairwise comparisons, if all these parametric assumptions were met, groups were compared using unpaired t-tests; if the groups were normally distributed but had non-homogeneous variance, comparisons were made using unpaired t-tests with Welch’s correction for unequal variances; if none of the parametric assumptions were met, groups were compared using Mann-Whitney U tests. After testing, p-values were adjusted using Bonferroni’s correction any time two or more pairwise comparisons were performed. The sample size for each comparison is indicated in each plot, and p-values, sample sizes, and parametric assumptions are reported in Supplementary Table 1.

## SUPPLEMENTARY MOVIE LEGENDS

**Movie S1. Example instance of a non-mating female displaying three consecutive bouts of aggressive behaviour towards the mating pair. Related to Figure 1 and Figure 3**. Each bout is identified by a co-occurring red circle labelled “Aggression” around the flies.

**Movie S2. Example instance of a non-mating female displaying three consecutive bouts of aggressive behaviour towards the mating pair in a new, more spatially constrained arena. Related to Figure 2 and Figure 5**. Each bout is identified by a co-occurring red circle labelled “Aggression” around the flies.

**Movie S3. Example instance of a non-mating female separated from the mating pair by a nylon mesh. Related to Figure 2 and Figure 5**. The mesh is present since the start of the experiment, when flies are introduced to the arena, and removed after 30 minutes, thus allowing for courtship and mating of the second, isolated female.

