## Supplementary material for "Mate Competition Drives Aggressive Behaviour in Female *Drosophila*": Key Resources Table

| REAGENT or RESOURCE | SOURCE | IDENTIFIER |
|---|---|---|
| Deposited data | | |
| Raw data and code | This paper | https://doi.org/10.7910/DVN/URA528 |
| Experimental models: Organisms/strains | | |
| Wild type (DL strain) | [78] | N/A |
| $w^{1118}$ ; Poxn$\Delta^{[M22\text{-}B5]}$ ; $\Delta^{[SfoBs105]}$ | [79] | N/A |
| $w^{1118}$ ; Poxn$\Delta^{[M22\text{-}B5]}$ ; $\Delta^{[SfoBs127]}$ | [79] | N/A |
| w[*], norpA$^{36}$ ; + ; + | [80] | RRID: BDSC_9048 |
| IR8a$^{1}$; IR25a$^{2}$; Orco$^{1}$, GR63a$^{1}$ | [81] | N/A |
| w[*] ; UAS-Kir2.1.EGFP ; + | [73] | RRID: BDSC_6596 |
| w[*] ; IR84a-GAL4 ; TM2/TM6b, Tb$^{1}$ | [82] | RRID: BDSC_41734 |
| w[*], OR67d-GAL4 ; + ; + | [83] | RRID: BDSC_9997 |
| w[*] ; OR47b-GAL4; + | [84] | RRID: BDSC_9983 |
| Software and algorithms | | |
| Caltech Flytracker | [77] | http://www.vision.caltech.edu/Tools/FlyTracker/ |
| Bonsai | [76] | https://edspace.american.edu/openbehavior/project/bonsai/ |
| PythonVideoAnnotator | https://biodata.pt/python_video_annotator | https://pythonvideoannotator.readthedocs.io/en/master/ |
| Python | http://www.python.org/ | http://www.python.org/ |
