## Supplemental Figures and Table for "Mate Competition Drives Aggressive Behaviour in Female *Drosophila*"

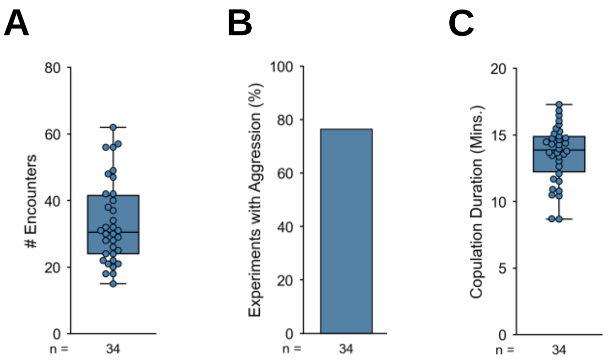


**Figure S1 | Characterization of female aggressive behaviour in the presence of a mating pair.**
**(A)** Number of encounters occurring during copulation. Encounters are defined as moments where the female is less than 4mm away from the mating pair.
**(B)** Aggression occurrence, expressed as the percentage of total experiments where aggression is observed.
**(C)** Duration, in minutes, of the first copulation (target female).

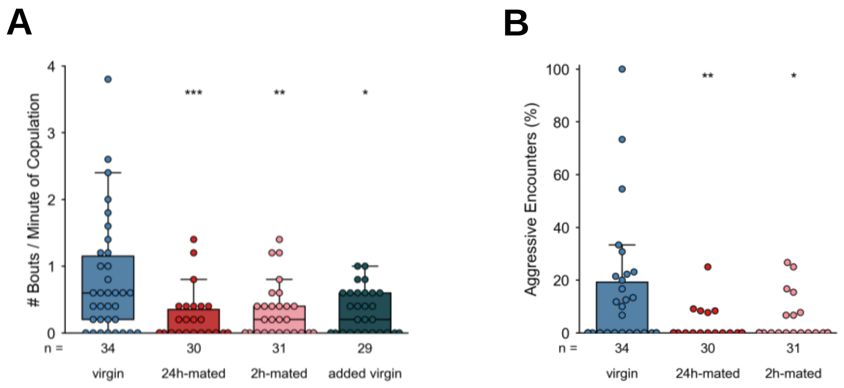


**Figure S2 | Aggression intensity is reduced by mating or lack of courtship exposure.**
**(A)** Number of aggression bouts per minute of copulation displayed by wild type virgin, 24h-mated, 2h-mated, and added virgin females towards a mating pair.
**(B)** Percentage of encounters where aggression occurs.
See Supplementary Table 3 for p-values and effect sizes; ns = not significant, *p < 0.05, **p < 0.01, ***p < 0.001. Sample size (number of flies tested) is shown for each condition below its corresponding boxplot.

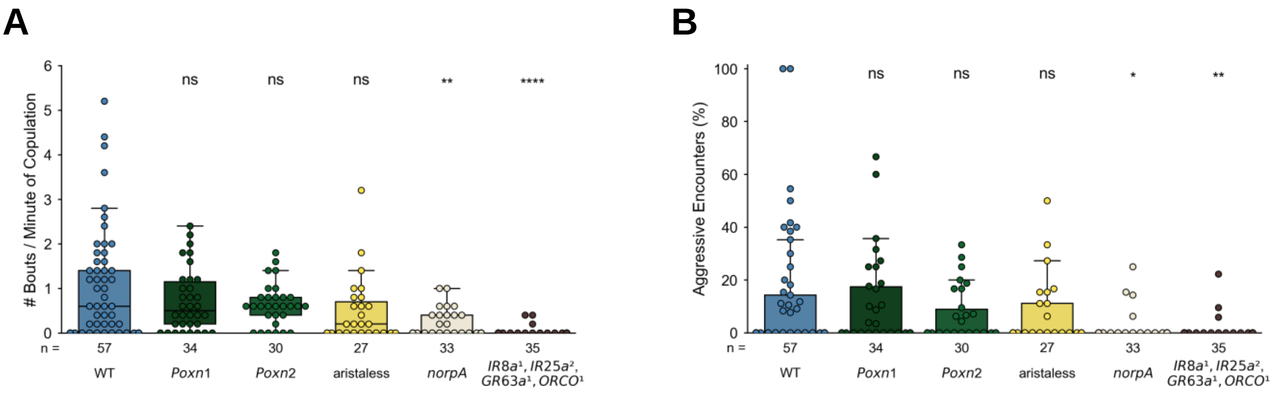


**Figure S3 | Olfaction and vision play an important role in female aggressive displays.**

**(A)** Number of aggression bouts per minute of copulation displayed by wild type, tasteless, deaf, blind, and anosmic virgin females towards a mating pair.

**(B)** Percentage of encounters where aggression occurs.

See Supplementary Table 2 for fly genotypes, and Supplementary Table 3 for p-values and effect sizes; ns = not significant, *p < 0.05, **p < 0.01, ****p < 0.0001. Sample size (number of flies tested) is shown for each condition below its corresponding boxplot.

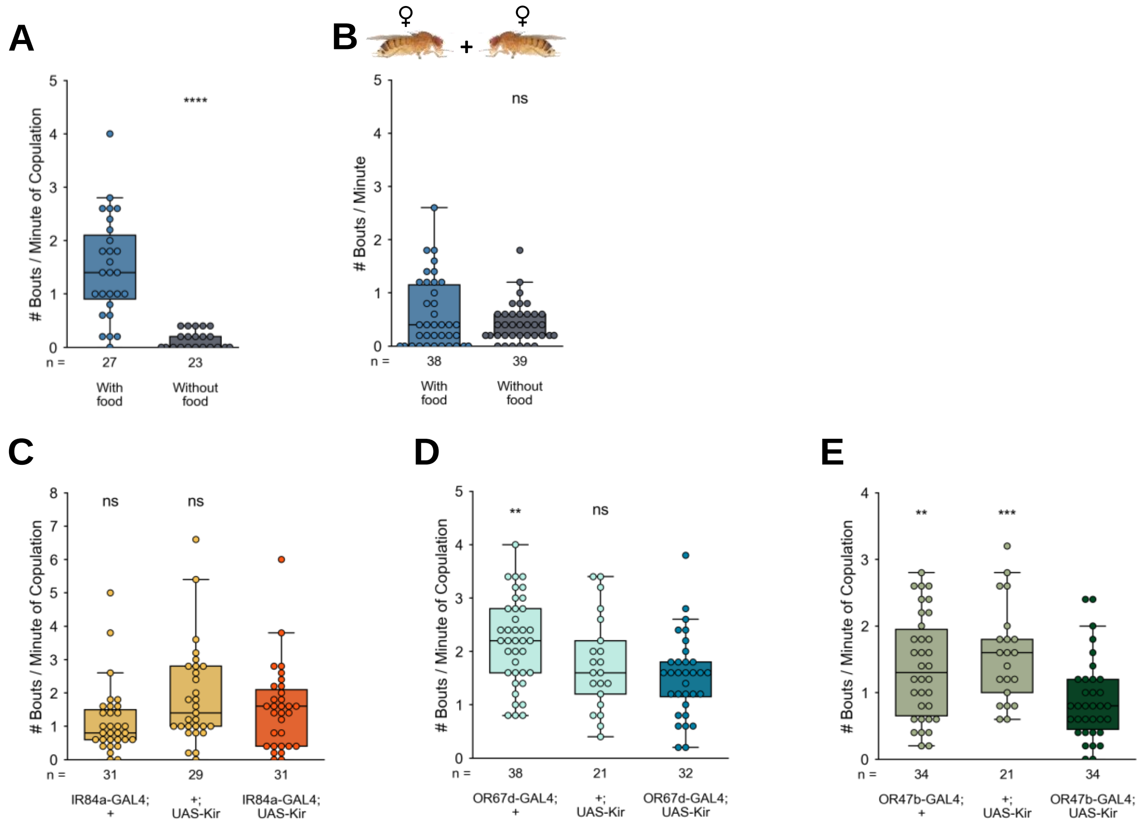


**Figure S4 | Presence of food odour and OR47b activity play an important role in female aggressive displays.**
Number of aggression bouts per minute in the following conditions:
**(A)** Wild type flies in the absence or presence of food odour.
**(B)** Wild type female pairs, without a male, in the absence or presence of food odour.
**(C)** Silenced IR84a flies and respective controls in the presence of food odour.
**(D)** Silenced OR67d flies and respective controls in the presence of food odour.
**(E)** Silenced OR47b flies and respective controls in the presence of food odour.
See Supplementary Table 2 for fly genotypes, and Supplementary Table 3 for p-values and effect sizes; ns = not significant, $**p < 0.01$, $***p < 0.001$, $****p < 0.0001$. Sample size (number of flies tested) is shown for each condition below its corresponding boxplot.

### Supplementary Table 1. Statistical details related to main and supplementary figures

| Figure | Groups | Sample size | Normally distributed[a] | Equal variance[b] | Statistical test | p-value | Effect size[c] |
|---|---|---|---|---|---|---|---|
| 2a | a) wild type female with aggression<br>b) wild type female without aggression (partition) | 34<br>28 | no<br>no | no | Mann-Whitney test | 0.566732 (ns) | 0.026 |
| | a) wild type female with aggression<br>c) wild type female without aggression (couple) | 34<br>40 | no<br>no | yes | Mann-Whitney test | 0.033061 (*) | 0.176 |
| 2b | a) wild type female with aggression<br>b) wild type female without aggression (partition) | 34<br>28 | no<br>no | yes | Mann-Whitney test | 0.139606 (ns) | - 0.107 |
| 2c | a) wild type female with aggression<br>b) wild type female without aggression (partition) | 34<br>28 | yes<br>yes | yes | Two sample t-test | 0.178776 (ns) | - 0.319 |
| | a) wild type female with aggression<br>c) wild type female without aggression (couple) | 34<br>40 | yes<br>no | yes | Mann-Whitney test | 0.591032 (ns) | - 0.064 |
| 2d | a) wild type female with aggression<br>b) wild type female without aggression (partition) | 34<br>28 | yes<br>yes | yes | Two sample t-test | 1.0 (ns) | 0.130 |
| | a) wild type female with aggression<br>c) wild type female without aggression (couple) | 34<br>40 | yes<br>no | yes | Mann-Whitney test | 0.428794 (ns) | 0.034 |
| 2e | a) wild type female with aggression<br>b) wild type female without aggression (partition) | 34<br>28 | no<br>yes | yes | Mann-Whitney test | 0.391206 (ns) | 0.146 |
| 2f | a) wild type female with aggression<br>b) wild type female without aggression (partition) | 34<br>28 | yes<br>yes | yes | Two sample t-test | 0.749854 (ns) | - 0.021 |
| 3a | a) wild type virgin female<br>b) wild type 24h-mated female | 34<br>30 | no<br>no | no | Mann-Whitney test | 0.000621 (***) | - 1.0 |
| | a) wild type virgin female<br>c) wild type 2h-mated female | 34<br>31 | no<br>no | no | Mann-Whitney test | 0.001358 (**) | - 0.950 |
| | a) wild type virgin female<br>d) wild type added virgin female | 34<br>29 | no<br>no | no | Mann-Whitney test | 0.003371 (**) | - 0.888 |
| 3b | a) wild type virgin female<br>b) wild type 24h-mated female | 34<br>30 | yes<br>no | yes | Mann-Whitney test | 0.329374 (ns) | 0.134 |
| | a) wild type virgin female<br>c) wild type 2h-mated female | 34<br>31 | yes<br>no | yes | Mann-Whitney test | 0.879943 (ns) | - 0.071 |
| 4a | a) wild type virgin female<br>b) tasteless mutant 1 virgin female | 57<br>34 | no<br>no | yes | Mann-Whitney test | 1.0 (ns) | - 0.118 |
| | a) wild type virgin female<br>b) tasteless mutant 2 virgin female | 57<br>30 | no<br>no | yes | Mann-Whitney test | 1.0 (ns) | 0.335 |
| | a) wild type virgin female<br>b) deaf virgin female | 57<br>27 | no<br>no | no | Mann-Whitney test | 0.178375 (ns) | - 0.757 |
| | a) wild type virgin female<br>b) blind mutant virgin female | 57<br>33 | no<br>no | no | Mann-Whitney test | 0.005105 (**) | - 1.0 |
| | a) wild type virgin female<br>b) anosmic mutant virgin female | 57<br>35 | no<br>no | no | Mann-Whitney test | $1.869\times10^{-7}$ (****) | - 1.0 |
| 4b | a) wild type virgin female<br>b) tasteless mutant 1 virgin female | 57<br>34 | yes<br>yes | yes | Two sample t-test | 1.0 (ns) | - 0.076 |
| | a) wild type virgin female<br>b) tasteless mutant 2 virgin female | 57<br>30 | no<br>no | yes | Mann-Whitney test | 1.0 (ns) | - 0.054 |
| | a) wild type virgin female<br>b) deaf virgin female | 57<br>27 | yes<br>yes | yes | Two sample t-test | 1.0 (ns) | - 0.148 |
| | a) wild type virgin female<br>b) blind mutant virgin female | 57<br>33 | yes<br>yes | yes | Two sample t-test | 1.0 (ns) | - 0.186 |
| | a) wild type virgin female<br>b) anosmic mutant virgin female | 57<br>35 | yes<br>yes | yes | Two sample t-test | 1.0 (ns) | - 0.009 |
| 5a | a) wild type virgin female (with food)<br>b) wild type virgin female (without food) | 27<br>23 | no<br>no | no | Mann-Whitney test | $2.941\times10^{-8}$ (****) | - 1.0 |
| 5b | a) wild type virgin female (with food)<br>b) wild type virgin female (without food) | 38<br>39 | no<br>no | no | Mann-Whitney test | 0.247603 (ns) | 0.886 |
| 5c | a) IR84a > Kir2.1 silencing<br>b) Gal4 control | 31<br>31 | no<br>no | yes | Mann-Whitney test | 0.159089 (ns) | 0.859 |
| | a) IR84a > Kir2.1 silencing<br>b) UAS control | 31<br>29 | no<br>no | yes | Mann-Whitney test | 0.700511 (ns) | - 0.207 |
| 5d | a) OR67d > Kir2.1 silencing<br>b) Gal4 control | 32<br>38 | no<br>yes | yes | Mann-Whitney test | 0.074072 (ns) | - 0.395 |
| | a) OR67d > Kir2.1 silencing<br>b) UAS control | 32<br>21 | no<br>no | yes | Mann-Whitney test | 0.591601 (ns) | - 0.241 |
| 5e | a) OR47b > Kir2.1 silencing<br>b) Gal4 control | 34<br>34 | no<br>no | yes | Mann-Whitney test | 0.037607 (*) | - 0.415 |
| | a) OR47b > Kir2.1 silencing<br>b) UAS control | 34<br>21 | no<br>no | yes | Mann-Whitney test | 0.014572 (*) | - 0.398 |
| S2a | a) wild type virgin female<br>b) wild type 24h-mated female | 34<br>30 | no<br>no | no | Mann-Whitney test | 0.000663 (***) | - 1.0 |
| | a) wild type virgin female<br>c) wild type 2h-mated female | 34<br>31 | no<br>no | no | Mann-Whitney test | 0.009462 (**) | - 0.667 |
| | a) wild type virgin female<br>d) wild type added virgin female | 34<br>29 | no<br>no | no | Mann-Whitney test | 0.049136 (*) | - 0.667 |
| S2b | a) wild type virgin female<br>b) wild type 24h-mated female | 34<br>30 | no<br>no | no | Mann-Whitney test | 0.007610 (**) | NA |
| | a) wild type virgin female<br>c) wild type 2h-mated female | 34<br>31 | no<br>no | no | Mann-Whitney test | 0.038226 (*) | NA |
| S3a | a) wild type virgin female<br>b) tasteless mutant 1 virgin female | 57<br>34 | no<br>no | yes | Mann-Whitney test | 1.0 (ns) | - 0.167 |
| | a) wild type virgin female<br>b) tasteless mutant 2 virgin female | 57<br>30 | no<br>no | no | Mann-Whitney test | 1.0 (ns) | 0.0 |

| | | | | | | | |
|---|---|---|---|---|---|---|---|
| | a) wild type virgin female<br>b) deaf virgin female | 57<br>27 | no<br>no | yes | Mann-Whitney test | 0.236137 (ns) | - 0.667 |
| | a) wild type virgin female<br>b) blind mutant virgin female | 57<br>33 | no<br>no | no | Mann-Whitney test | 0.001278 (***) | - 1.0 |
| | a) wild type virgin female<br>b) anosmic mutant virgin female | 57<br>35 | no<br>no | no | Mann-Whitney test | $2.247 \times 10^{-7}$ (****) | - 1.0 |
| S3b | a) wild type virgin female<br>b) tasteless mutant 1 virgin female | 57<br>34 | no<br>no | yes | Mann-Whitney test | 1.0 (ns) | NA |
| | a) wild type virgin female<br>b) tasteless mutant 2 virgin female | 57<br>30 | no<br>no | yes | Mann-Whitney test | 1.0 (ns) | NA |
| | a) wild type virgin female<br>b) deaf virgin female | 57<br>27 | no<br>no | yes | Mann-Whitney test | 1.0 (ns) | NA |
| | a) wild type virgin female<br>b) blind mutant virgin female | 57<br>33 | no<br>no | no | Mann-Whitney test | 0.020358 (*) | NA |
| | a) wild type virgin female<br>b) anosmic mutant virgin female | 57<br>35 | no<br>no | no | Mann-Whitney test | 0.004451 (**) | NA |
| S4a | a) wild type virgin female (with food)<br>b) wild type virgin female (without food) | 27<br>23 | yes<br>no | no | Mann-Whitney test | $4.357 \times 10^{-8}$ (****) | - 1.0 |
| S4b | a) wild type virgin female (with food)<br>b) wild type virgin female (without food) | 38<br>39 | no<br>no | yes | Mann-Whitney test | 0.419892 (ns) | 0.0 |
| S4c | a) IR84a > Kir2.1 silencing<br>b) Gal4 control | 31<br>31 | no<br>no | yes | Mann-Whitney test | 0.172588 (ns) | 1.0 |
| | a) IR84a > Kir2.1 silencing<br>b) UAS control | 31<br>29 | no<br>no | yes | Mann-Whitney test | 0.410530 (ns) | 0.143 |
| S4d | a) OR67d > Kir2.1 silencing<br>b) Gal4 control | 32<br>38 | yes<br>yes | yes | Two sample t-test | 0.001858 (**) | - 0.273 |
| | a) OR67d > Kir2.1 silencing<br>b) UAS control | 32<br>21 | yes<br>yes | yes | Two sample t-test | 0.544273 (ns) | 0.0 |
| S4e | a) OR47b > Kir2.1 silencing<br>b) Gal4 control | 34<br>34 | no<br>yes | no | Mann-Whitney test | 0.008534 (**) | - 0.385 |
| | a) OR47b > Kir2.1 silencing<br>b) UAS control | 34<br>21 | no<br>yes | yes | Mann-Whitney test | 0.000599 (***) | - 0.5 |

ns = not significant
* = p-value < 0.05
** = p-value < 0.01
*** = p-value < 0.001
**** = p-value < 0.0001
[a] To test for normal distribution: Shapiro's test and D'Agostino's test.
[b] To test for the homogeneity of variance: Levene´s test for non-normally distributed samples and Bartlett´s test for normally distributed samples.
[c] To calculate effect size: Median fold change = (test group median - control group median) / control group median.
